# ANTIPASTI: interpretable prediction of antibody binding affinity exploiting Normal Modes and Deep Learning

**DOI:** 10.1101/2023.12.22.572853

**Authors:** Kevin Michalewicz, Mauricio Barahona, Barbara Bravi

**Affiliations:** Department of Mathematics, Imperial College London, London SW7 2AZ, United Kingdom

**Keywords:** Antibody, Binding Affinity, Deep Learning, Interpretability, Normal Mode Analysis, Protein Structures

## Abstract

The high binding affinity of antibodies towards their cognate targets is key to eliciting effective immune responses, as well as to the use of antibodies as research and therapeutic tools. Here, we propose ANTIPASTI, a Convolutional Neural Network model that achieves state-of-the-art performance in the prediction of antibody binding affinity using as input a representation of antibody-antigen structures in terms of Normal Mode correlation maps derived from Elastic Network Models. This representation captures not only structural features but energetic patterns of local and global residue fluctuations. The learnt representations are interpretable: they reveal similarities of binding patterns among antibodies targeting the same antigen type, and can be used to quantify the importance of antibody regions contributing to binding affinity. Our results show the importance of the antigen imprint in the Normal Mode landscape, and the dominance of cooperative effects and long-range correlations between antibody regions to determine binding affinity.

## 1 Introduction

Antibodies are proteins that play a major role in the immune response by binding to harmful foreign agents, *i.e*., antigens. Antibodies are both highly effective and specific in their binding capabilities. These properties, which are crucial when targeting antigens responsible for infections, have also led to the development of engineered antibodies for the treatment of other diseases, *e.g*., by targeting cancer cells [1, 2, 3].

Structurally, antibodies have a prototypical Y-shape: the stem of the ‘Y’ is invariant and responsible for the communication with the rest of the immune system, whereas each of the two identical tips of the ‘Y’ comprises variable regions that are tailored to specific antigens [4]. Each tip is composed of a light and a heavy chain [5], which contain the entire *paratope, i.e*., the collection of antibody residues that interact with the antigen’s *epitope*. Most of the paratope residues are distributed in CDRs (complementarity-determining regions), three on each chain. In general, CDRs involve between 6 and 20 amino acids [6], with the CDR-H3 (third complementarity-determining region of the heavy chain) typically harbouring more epitope binding sites than other regions [7, 8, 9]. Antibodies that are not yet fully developed, and hence not specialised for a particular epitope, are called *germline* or *naïve*. Upon exposure of the immune system to an antigen, a process entailing somatic mutations leads to *matured* antibodies capable of binding with increased affinity to the specific target.

There is an increasing number of datasets of antibodies and their corresponding targets, which include 1 usually amino acid sequences [10, 11] and sometimes three-dimensional structures [12, 13, 14]. As such, they have enabled the training of Deep Learning models for function prediction and characterisation even from datasets of relatively modest size [15]. In general, Deep Learning addresses different tasks, such as regression or classification, by means of multiple non-linear layers organised according to particular architectures. In the case of antibody modelling, Convolutional Neural Networks (CNNs) have been applied to structural data to predict contacts [16] and antibody-antigen binding interfaces [17], and to find potential binders for epitopes of interest [18]. Convolutions are powerful when dealing with protein structures due to their robustness in detecting features, regardless of their exact position [19] through a consistent set of learnable parameters. This characteristic makes it possible to find common structural motifs, which can provide insights into protein function [20]. Furthermore, given the multi-scale nature of proteins, the hierarchical architecture of CNNs is ideal for capturing features at different levels of abstraction [21].

While less available as of yet, structural data are often more informative about the specific configuration of binding, and hence affinity. Indeed, sequence-based methods have proven to be less effective, with a reduced ability to capture spatial synergies, as shown by the modest reported accuracies in Refs. [22, 23]. To date, only a few methods have used structural information to predict antibody binding affinity [24, 25, 26]. However, these approaches do not use the entire variable region of the antibody, relying instead on a handcrafted engineering of a few features as paratope contact-based and area-based descriptors. Yet, given that protein structures are specified by hundreds to thousands of atomic spatial coordinates, methods trained on structures tend to be computationally expensive [27]. Ideally, one would want to find an embedding that captures the structural properties of all the amino acids in the antibody variable region as well as those of the antigen, while also taking into account their relationships. The choice of a parsimonious yet informative input data representation becomes particularly crucial to developing any such structure-based approaches. As described below, our proposed method uses CNNs applied to residue-level descriptions of antibodies to circumvent these problems.

The need to include broader structural information stems from the fact that the molecular determinants of binding affinity are not fully understood, thus affecting our ability to identify sites and regions to target for antibody engineering. For instance, although the CDR-H3 is generally acknowledged to be the main driver of high-affinity binding [28], several studies indicate that the other CDRs also play an important role [29, 30]. Furthermore, interactions across antibody regions are observed to favour high binding affinity, including bonds between the FR-L2 and heavy chain CDR loops, which help stabilise and orient them into the antigen-bound conformation [30].

In a different line of research, delivering generalisable insights on how structural rigidity affects binding affinity has also proved difficult. On one hand, a positive correlation between rigidity and high affinity has been reported [31, 32, 33, 34, 35] and increased localised rigidity with lower residue fluctuations (B-factors) has been found in the CDR-H3 of affinity-matured antibodies relative to their *naïve* counterparts [36]. On the other hand, it was found that maturation reduces flexibility only if there is initially a strong binding to the conserved antigen sites [37], and that mutations that stabilise CDR loops in a binding-compatible conformation can confer high binding affinity without CDR-H3 rigidification [38].

The computational study of such structural fluctuations via molecular dynamics is hampered by the long-time scales involved, with only a limited number of case studies available [37, 39, 40]. Alternatively, Normal Mode Analysis (NMA) [41] allows the study of correlated molecular fluctuations in protein structures, and is especially efficient when applied to coarse-grained descriptions of proteins such as Elastic Network Models (ENMs) [42, 43], which model the protein as a network of residues interacting via empirically calibrated harmonic potentials [44]. The Normal Modes of such networks capture fluctuations around equilibrium with different frequencies and spatial ranges. Although NMA is well established, with several software tools widely used by the community [45, 46, 47], it has not been exploited to characterise the structural fluctuations of antibody-antigen complexes.

In this work, we develop ANTIPASTI (ANTIbody Predictor of Affinity from STructural Information), an interpretable Deep Learning approach that harnesses the information encoded in the molecular structure and its associated interactions for the numerical prediction of binding affinity. Specifically, ANTIPASTI is based on a CNN architecture that learns a representation of antibody structures from binding affinity data and residue-level Normal Mode correlation maps derived from Elastic Network Models. The learnt representation can then be used to predict the binding affinity of any given structure and to extract interpretable structural features connected to antibody regions that are key contributors to increase and decrease binding affinity.

## 2 Results

### The ANTIPASTI Deep Learning architecture

ANTIPASTI uses structural data of antibody-antigen complexes together with experimentally measured values of their binding dissociation constant (*K*_*D*_) to train a Deep Learning model based on a CNN architecture in a supervised fashion. Our training dataset consists of 634 antibody-antigen structures, retrieved from SAbDab [12], annotated with an experimental *K*_*D*_ value (see Methods). The input to our model is a residue-residue correlation map. This map is calculated from the Normal Modes of an Elastic Network Model (ENM) of the antibody-antigen complex, a residue-level representation of the structure containing energy interactions (Methods). The Normal Mode Analysis of the ENM antibody-antigen structural complex encodes quantitative information on regions that undergo correlated or anti-correlated fluctuations (Figure 1a). Since binding affinity is determined by the specific conformation of the antibody and its cognate antigen, we compute the input correlation map of the *full* antibody-antigen complex, retaining in this way the antigen’s imprint on the antibody residues (Methods). We then extract the portion of the Normal Mode map corresponding to the antibody residues to obtain our input, which is set to have the same size by adopting a uniform numbering scheme for antibody residue positions (Chothia scheme [49], see Figure 1b).

**Figure 1:**
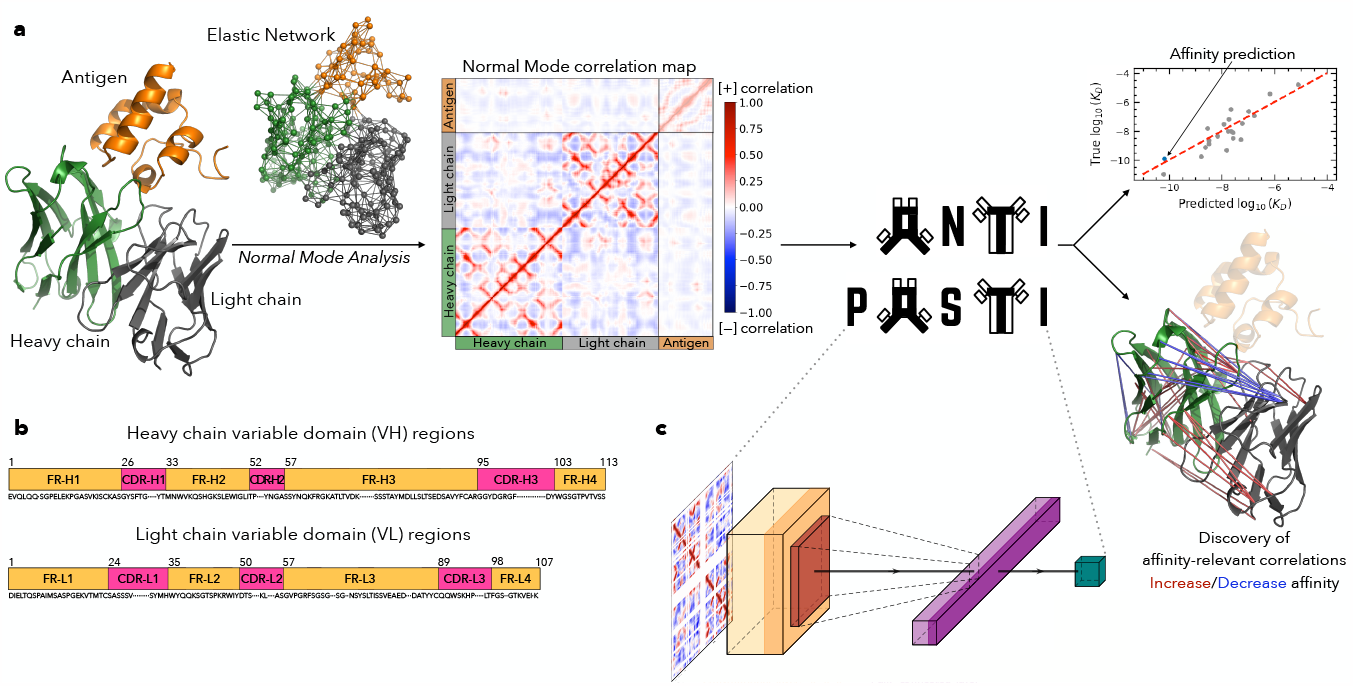
Overview of ANTIPASTI. (**a**) Schematic of the approach (structure image created with [48]). (**b**) Regions are defined with an alignment to the Chothia position numbering scheme [49]. (**c**) Architecture of the ANTIPASTI Deep Learning model (see Methods).

The ANTIPASTI neural network architecture is composed of a CNN with *n*_f_ filters of shape *k* × *k*, a non-linearity and a *p*-pooling layer, followed by a fully-connected layer, which outputs the log_10_(*K*_*D*_) prediction (Figure 1c, Methods). The representation of antibody structures via Normal Mode correlation maps provides image-like inputs, for which CNNs, firstly developed for applications in computer vision [19], are particularly well-suited. During training, the model learns new representations of antibody structures guided by the supervised task of mapping residue-residue correlations in the antigen-bound conformation to binding affinity. The representations are kept interpretable as they can be mapped back to the original antibody structure. To ensure reproducibility, our pipeline and data are publicly available with instructions and tutorials provided on GitHub (see Code availability).

Given structural data for an antibody-antigen complex, ANTIPASTI first computes the ENM representation and its corresponding Normal Mode correlation map for the antibody (under the imprint of the antigen). This correlation map is then used as the input to the CNN model to predict the binding affinity constant. In addition, the representations can be inspected to discover which correlations and antibody regions are most relevant to binding affinity (Figure 1c).

### ANTIPASTI accurately predicts binding affinity

We conducted hyperparameter optimisation and identified a CNN with pooling as the best ANTIPASTI architecture, which we will use in the following sections for various analyses (Figure S1). The second best model was a CNN without pooling, which we included in our accuracy tests. We evaluated these two best model candidates on the test set for five random seeded train-test splits (see Methods for more details) and we measured their performance via the *R* correlation score between the true *K*_*D*_ and the predicted values by ANTIPASTI. As a baseline, we also considered a basic linear regression to predict binding affinity from Normal Mode correlation maps (Methods).

We then investigated the accuracy of ANTIPASTI for prediction of the binding affinity constant *K*_*D*_. Figure 2a shows the predicted *K*_*D*_ values of the CNN with pooling on an unseen test set compared to the true values with a Pearson correlation *R* = 0.86. The mean *R* score computed over 5 test sets is *R* = 0.85. The other architecture (CNN without pooling) exhibits comparable performance, with mean *R* score of 0.86 (Figure 2c). Such a performance compares favourably with existing sequence-based and structure-based predictors of antibody binding affinity [23, 22, 24, 26], which have *R* scores ranging from 0.45 − 0.79, albeit a full comparison cannot be carried out consistently (Table S1). This is due to differences in method design and data selection, which prevent us from re-training and evaluating their models on our more general data. For comparison, the linear regression on Normal Modes achieves a mean *R* = 0.64, substantially lower than ANTIPASTI (Figure 2c), indicating that is crucial to include the non-linearities in the network layers with robust feature detection properties, such as convolutional layers. Furthermore, we note that the ANTIPASTI architectures are computationally efficient: the training time of all the architectures implemented here remained below 10 minutes (Apple M1, macOS Big Sur).

**Figure 2:**
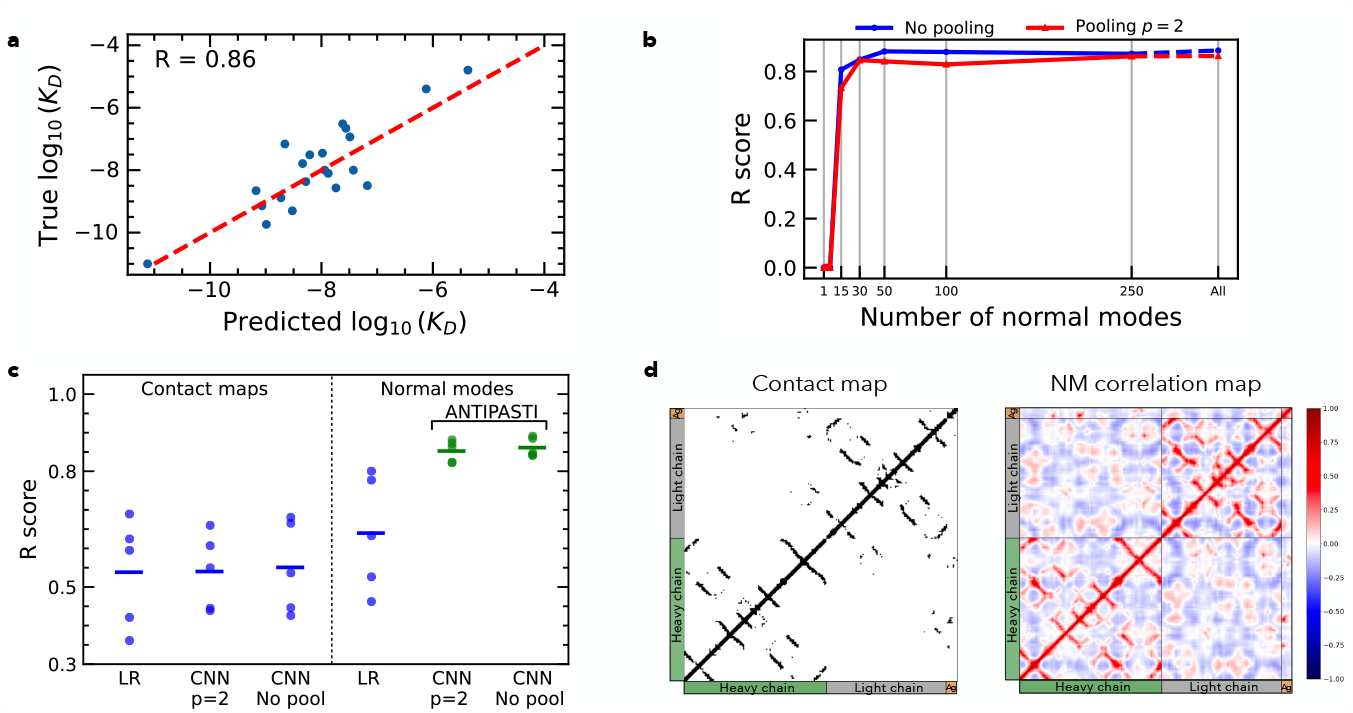
ANTIPASTI performance. (**a**) ANTIPASTI predictions for a particular test set that yielded a correlation coefficient *R* = 0.86 between true and predicted log_10_ (*K*_*D*_) values. (**b**) Test performance as a function of the number of Normal Modes used. Note that the first six are trivial and that they are ordered from the lowest to highest associated eigenvalue (Methods). (**c**) Test performance for five distinct train-test random splits comparing the two best ANTIPASTI architectures - with *p* = 2 pooling and no pooling - to simpler approaches: linear regression (LR), and CNNs and LR trained on contact maps. Mean *R* scores are indicated by a horizontal line. (**d**) Example of a contact map and Normal Mode correlation map (PDB entry: `3fn0`).

We next analysed the robustness of our results to the number of Normal Modes used to build the input map. Normal modes are ordered by frequency (from low to high), hence capturing from global to increasingly more local structural fluctuations. A natural question is if the number of modes could be reduced while maintaining the predictive performance or, in other words, what spatial scales of correlated fluctuations matter for binding affinity. We observed that the prediction accuracy remained high when excluding high frequency Normal Modes: as shown in Figure 2b, the performance of ANTIPASTI is unaffected as long as the first 30 modes (*i.e*., those with lowest frequency and largest spatial scales) are kept (*R* = 0.845). This indicates that binding affinity is predominantly affected by long-range spatial correlations, while short-range correlations can be discarded with little loss of prediction power for binding affinity.

### Normal Mode correlation maps encode richer information than contact maps

We also explored whether residue-residue contact maps, where a contact is recorded if the physical distance between *α*-Carbon atoms is less than a threshold (typically 6 − 12Å [50]), are sufficient to estimate *K*_*D*_ instead of opting for Normal Modes. To tackle this question, we carried out a hyperparameter optimisation for this type of input and then re-trained ANTIPASTI using contact maps as input data (Methods).

Using the same protocol as for our Normal Mode computations, we found that CNN models applied to contact map inputs lead to test set *R* scores ranging between 0.36 and 0.68 (Figure 2c). This confirms that the antibody structural connections mediated through 3D space are informative, yet substantially less so than the structural fluctuations (*dynamical* as much as structural information) captured by Normal Mode maps. Note also that linear regression and CNN models perform indistinguishably on contact map inputs (Figure 2c), a fact that can be explained due to the sparsity of contact maps (Figure 2d), *i.e*., the large amount of zero pixels makes the hierarchical convolutional coarse-graining less useful.

### ANTIPASTI representations capture distinctive binding modes for different antigen types

The internal representations learnt by the model during training (specifically the output layer) can be used to reveal pairwise residue correlations that are key to binding affinity. Note that the sum of the elements of the output layer gives log_10_ (*K*_*D*_) (see Methods). Yet the individual elements contain information about correlations that produce an increase or a decrease of the predicted binding affinity. To visualise this, we reshape the output layer into a residue-residue *affinity-relevant correlation map* (Figure 3a, Figures 3d-e, *Methods)*.

**Figure 3:**
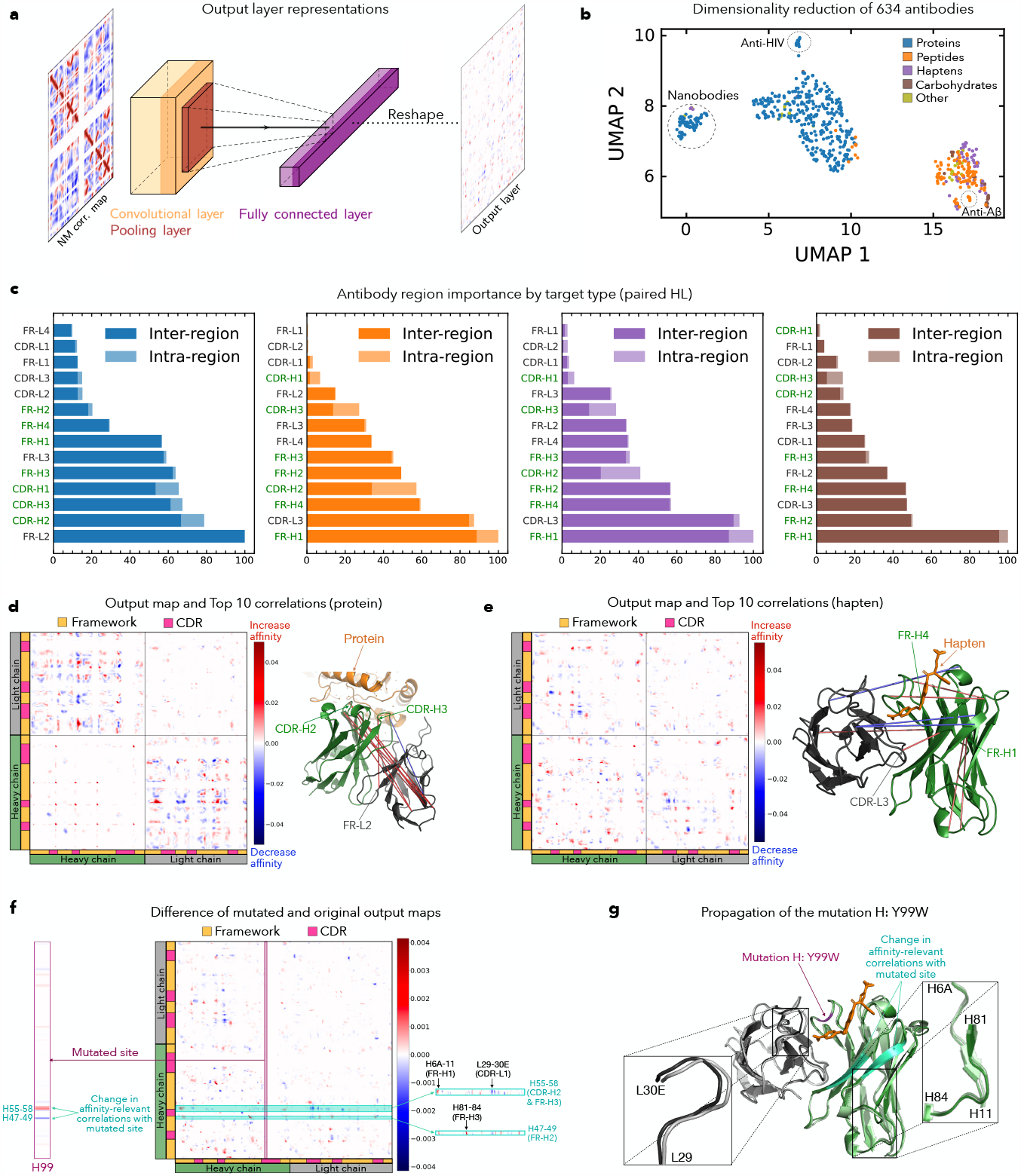
Residue-residue correlations relevant to binding affinity. (**a**) ANTIPASTI output representations indicating which correlations increase or decrease binding affinity. (**b**) Dimensionality reduction (UMAP) of the output layer representations groups antibodies by their binding target type. (**c**) Rankings of antibody regions’ importance, expressed as a percentage of the importance of the best region, for 4 the groups of antibodies with different target type highlighted in b (antibodies with paired heavy and light chains only). In general, inter-region affinity-relevant correlations contribute more to importance than intra-region ones. (**d**) Relevant correlations for binding affinity for an antibody with a protein target (Mntc with Mab 305-78-7 complex, PDB entry: `5hdq`). (**e**) Relevant correlations for binding affinity in the complex of an antibody targeting the hapten dabigatran (PDB entry: `4yhi`). (**f**) Difference in affinity-relevant correlations between a mutated antibody (`4yho`) and its original structure (`4yhi`), whose variable regions differ only at site H99 (CDR-H3). The mutated antibody presents a higher affinity. (**g**) `4yhi` and `4yho` structure overlay. The greatest variations in affinity-relevant correlations with the mutated site identify two *β*-strands (green-cyan). These are in turn correlated with numerous sites, three of which correspond to the most substantial conformational shifts between `4yhi` and `4yho`.

To further understand the patterns of affinity-relevant correlations, we applied UMAP dimensionality reduction [51] to the output layer representations of all available antibodies in our dataset. We found that antibodies cluster according to the type of antigen in the UMAP projection (Figure 3b) even though the inputs are restricted only to antibody residues. This finding confirms that the model leverages similarities in the pattern of correlated fluctuations induced by the imprint of antigens. In particular, antibodies binding to small size linear targets (*i.e*., haptens and peptides) cluster closely, hinting that target size and the presence of a secondary structure in the antigen impose constraints on binding. On the other hand, nanobodies stand out as a separate cluster, in line with the biologically sensible expectation that a distinctive binding conformation characterises antibodies with a single domain. Similarly, there is a tight cluster of anti-HIV antibodies comprising 14 structures, all of which correspond to the same species (Homo sapiens); have the same type of light chain (*κ*); and belong to the same heavy chain V gene family (IGHV1). Another tight cluster is seen for 6 antibodies targeting the Amyloid Beta peptide (Alzheimer’s disease), also sharing these properties in most cases (Mus musculus/*κ*/Other) (Figure 3b, Figure S3).

### ANTIPASTI representations identify important regions for binding affinity

We next used the ANTIPASTI output layer representations to define a measure of antibody region importance towards high binding affinity. Our measure quantifies the loss in accuracy due to the exclusion of the correlations of the residues in a particular region with all other sites, normalised to be a percentage of the top region (Methods). The results of the region importance measure are presented in Figure 3c for the antibodies grouped by antigen type.

As a general trend, antibodies that have haptens and peptides as antigens exhibit a similar ranking of important regions, different to antibodies targeting carbohydrates and especially proteins. This is consistent with the separation of protein-targeting antibodies in the UMAP projection (Figure 3b). In addition, heavy chain regions have high importance: 5 (4 for carbohydrates) of the 7 most relevant regions belong to the heavy chain. In particular, the CDR loops of the heavy chain are among the first 4 regions for antibodies with protein targets. Indeed, heavy chain CDR loops are known to harbour the majority of epitope-binding sites [52, 8] some of which have prominent effects on affinity to protein targets when mutated [52, 38]. CDR loops are surpassed only by the second framework of the light chain (FR-L2). Previous experimental work has shown that mutations in FR-L2 can lead to improved thermostability, a pre-requisite to binding [52], and residues within FR-L2 are part of the heavy-light chain interface involved in highly stabilising interactions with the CDR-H3, thus modulating its rigidity [32, 38] (see Figure 3d for an example showing highly important correlations linking CDR-H3 and FR-L2). Framework regions, in particular FR-H1,2,4, also stand out in the importance ranking for smaller targets (Figure 3c), consistent with an active role of framework sites in affinity and stability by influencing conformational dynamics during antigen binding [52].

Altogether, these observations suggest synergistic effects between regions in determining high-affinity binding. To quantify this further, we determined the proportion of importance due to the inter-region *vs*. intra-region correlations. We found that higher importance of inter-region correlations, underlining the role of synergy between regions in antigen binding. Repeating this computation to quantify intra-chain *vs*. inter-chain correlations corroborated that the importance of light chain framework regions stems to a large extent from correlations with the heavy chain regions (Figure S4a).

### ANTIPASTI representations reveal long-range correlations as key to affinity

To further validate the insight that affinity-relevant correlations span large distances in the structure, we considered the 10 most important correlations for each structure in the dataset. We found that most of them are long-range: 80% of the top interactions are between residues more than 10Å apart. The distribution also has a peak at short-distance, indicating that the model detects important short-range physical interactions (Figure S5).

To better understand affinity-relevant long-range correlations predicted by ANTIPASTI, we sought experimental data on single site mutations with an effect on antibody affinity. We could only find one mutational study [53] reporting both affinity values and the corresponding antigen-antibody complex structure, corresponding to antibody fragment idarucizumab targeting the hapten dabigatran. The original complex (PDB code: `4yhi`, Figure 3e) had *K*_*D*_ = 180 *pM*, and a single site mutation within the CDR-H3, from tyrosine to tryptophan (mutation H:Y99W), led to a 10-fold improvement in binding affinity in the mutated complex (PDB code: `4yho`). Although globally similar, there are noticeable changes in the affinity-relevant correlations following the mutation centered around two *β*-strands (H47-49 and H55-58, Figure 3f), which partly lie at the interface between the heavy and light chains (Figure 3g). In particular, H47-49 belongs to the binding pocket of dabigatran. Via structural analyses, Ref. [53] attributed the increased binding affinity to an optimisation of the steric fit to the paratope due to the incorporated tryptophan in the CDR-H3 pushing dabigatran towards the side of the binding pocket (residue H29 and *β*-strand residue H49), forming a new *π*-stacking bond. Indeed, the two *β*-strands (H47-49 and H55-58) undergo conformational shifts following the mutation (see zoomed insets on the overlaid structures in Figure 3g) and the loss of a hydrogen bond between L30C and H99 upon the mutation might lead to a tighter fit of dabigatran towards H29/H49 [53]. These changes lead to large differences in the ANTIPASTI-predicted affinity-relevant correlations with two neighbouring *β*-strands, which amplify and mediate the spread of the mutational effect to more flexible regions in the antibody (Figure 3f). Note that these two *β*-strands already appear in the top 10 correlations predicted by ANTIPASTI for the original (unmutated) structure (Figure 3e), as they present correlations with the CDR-L1 and CDR-L3, thus hinting at their potentially central role for modifying binding affinity.

### Evaluation tests with AlphaFold-predicted structures

Finally, we assessed the accuracy of ANTIPASTI affinity predictions for AlphaFold-predicted structures. Predicted structures are increasingly used as tools to understand protein functions, for homology detection [54] and to study the impact of point mutations [55]. To determine whether the performance of ANTIPASTI on AlphaFold predicted structures is comparable to experimental PDB structures, we generated AlphaFold structures [56, 57] from the sequences of the 21 antibody-antigen complexes of one test set (Methods). We then computed the Normal Mode correlation maps of these predicted structures and used them as inputs to ANTIPASTI to predict their binding affinity, 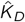. We found that the affinity prediction is comparable to the original PDB structures (Pearson’s correlation coefficient *R* = 0.97, Figure 4a), and the retrieval of the ground truth affinity *K*_*D*_ is unaffected (*R* = 0.84 *vs. R* = 0.86, see Figure S6a).

**Figure 4:**
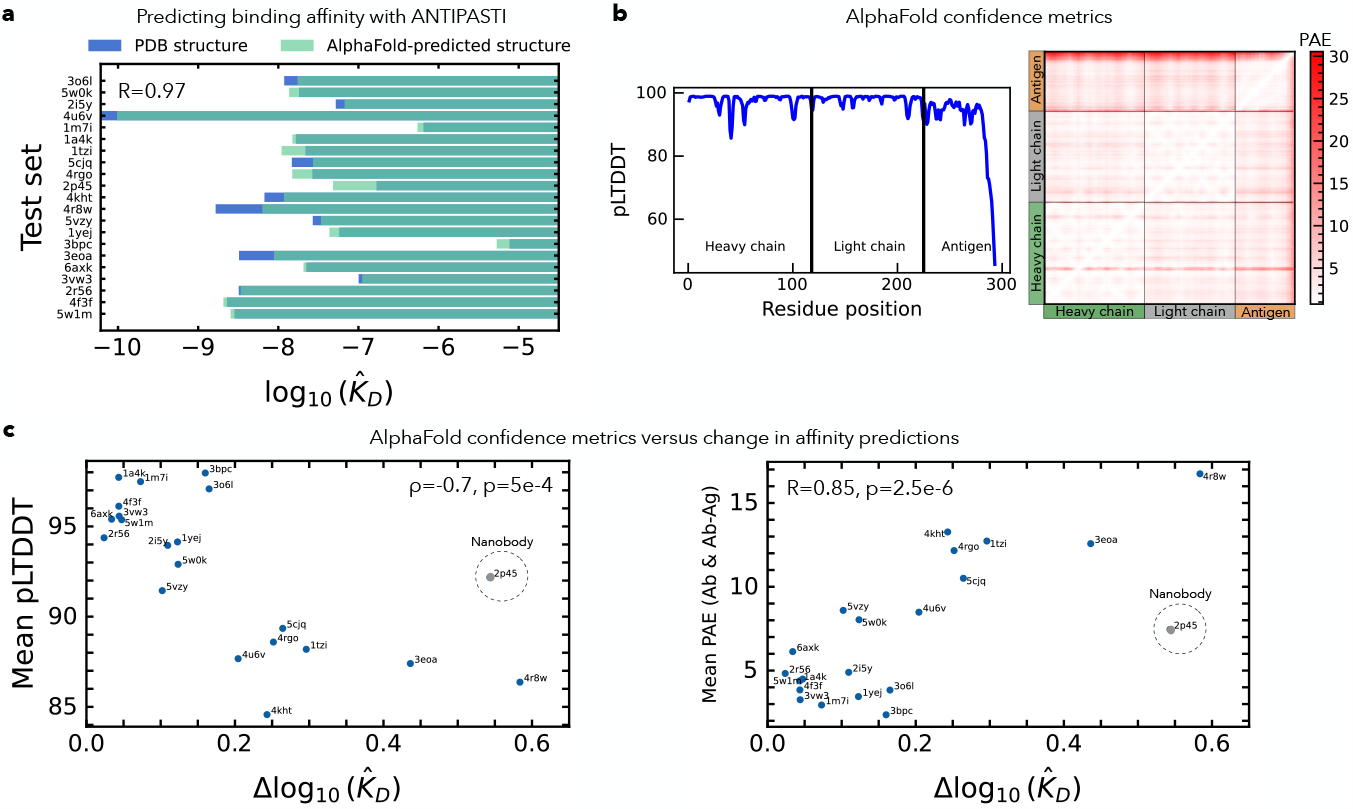
Evaluation of ANTIPASTI on AlphaFold-predicted structures. (**a**) Comparison of log_10_(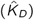) between the PDB structures and those predicted by AlphaFold for the same test set as in Figure 2a. (**b**) Alphafold confidence in folding prediction for the Msln7-64 Morab-009 Fab complex (PDB entry `4f3f`). The predicted Local Distance Difference Test (pLDDT) for each residue and the Predicted Alignment Error (PAE) matrix are displayed. (**c**) Mean of the per-residue confidence, assessed by the pLDDT metric (Spearman *ρ* = *−*0.7), and the mean PAE for the antibody residues and the antibody-antigen off-blocks (Pearson *R* = 0.85), as functions of Δlog_10_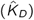.

To quantify further the effect of the uncertainty in AlphaFold predictions, we used the two AlphaFold-provided confidence metrics: the predicted Local Distance Difference Test (pLDDT) for each residue, and the Predicted Alignment Error (PAE) matrix. Figure 4b shows these metrics for the predicted complex with PDB entry `4f3f`. Except for the last residues of the antigen, the per-residue confidence is greater than 85 for all residues and the maximum PAE does not exceed 15. This translates into a low difference between the ANTIPASTI estimate of binding affinity for the original PDB structure and the Alphafold-predicted structure (Δlog_10_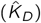) of less than 0.1 (Methods).

As expected, the larger the uncertainty in the AlphaFold prediction the larger the difference between the predicted affinities for experimental and AlphaFold structures. This can be seen in the negative correlation when plotting the average pLTDDT as a function of Δlog_10_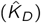 (Figure 4c, Spearman’s correlation coefficient *ρ* = −0.7 with a p-value of 5*e* − 4). Although there is also a positive correlation between the mean PAE versus Δlog_10_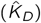 when considering the antibody block of the PAE matrix (Figure S6b, Pearson’s coefficient *R* = 0.74), the correlation increases to *R* = 0.85 (Figure 4c) when adding the antibody-antigen blocks, *i.e*., the AlphaFold uncertainty located at the interface between antibody and antigen residues. This result indicates the need for accurate AlphaFold prediction of the antibody-antigen interface to obtain a consistent affinity estimation with ANTIPASTI.

## 3 Discussion

In this work, we have developed ANTIPASTI, a new Machine Learning tool that leverages structural fluctuation information (obtained from Normal Mode correlation maps) to predict the binding affinity of antibodies using a CNN with a single filter bank and a fully-connected layer. We showed that ANTIPASTI can predict *K*_*D*_ from Normal Mode correlation maps with high generalisation power and achieves greater accuracy than existing methods (test set *R* = 0.86). Furthermore, training and inference are possible with low computational cost and using only a CPU.

A main ingredient of our approach is the use of residue-residue correlation maps obtained through Normal Mode Analysis of Elastic Network Model representations of antibody-antigen complexes as inputs that capture structural correlations without the need for long molecular dynamics simulations. Although the antigen is part of the NMA analysis, so as to retain the antigen imprint in the Normal Mode landscape of the antibody, the model is then trained on correlation maps of the imprinted antibody section alone. This choice allows ANTIPASTI to focus on the properties and regions of antibodies determining optimal binding, and can also potentially widen its scope of application to data that lack the cognate antigen in the resolved structure. To examine the importance of the antigen imprint for our method, we also tested an antigen-agnostic version (where the NMA analysis is carried out without the antigen section), and we found reduced predictive power, but still with information on biological characteristics in the output layer maps (Figure S2). The slow and large-scale fluctuations linked to low frequency modes are found to be most relevant to the prediction of binding affinity, suggesting long-range effects of antigen-binding conformational changes, possibly mediated through allosteric mechanisms [58, 59]. An aspect worth further investigation is the relative advantages of coarse-grained *vs*. atomistic protein structure representations [60].

We have also assessed the impact on the estimated binding affinity of AlphaFold-predicted, as opposed to experimentally resolved, structures. Our results on a small test set showed that evaluating a model trained on solved structures performs equally well on both types of structures giving support to the application of ANTIPASTI to predict the binding affinity of antibodies whose structure has not yet been experimentally obtained. We also found that the the discrepancy with the affinity estimated from AlphaFold structures (relative to the one estimated from the original structure) increases with the uncertainty of the structural prediction. Hence the AlphaFold-provided measure of uncertainty can be used as an indication on the level of certainty of the prediction. On a more conceptual level, our analysis indicated that the accurate prediction of the antibody-antigen contact interface is crucial to improve AlphaFold-predicted structures for the purpose of affinity prediction. We remark, that we have not used AlphaFold-predicted structures to augment the ANTIPASTI training set, as this can pose problems of data quality because at present AlphaFold is not guaranteed to predict accurately antibody-antigen complex structures [57, 61].

The learnt CNN filters can be used to establish which antibody regions are important for binding affinity. We find differences according to the type of antigen, especially between proteins and other small targets that presumably reflect patterns of binding configuration induced by the size of the target. Our model also allows us to establish an intrinsic measure of region importance based on to their impact on affinity prediction. Further work to establish model-agnostic approaches to quantifying importance through input perturbation [62], or occlusion sensitivity [63] would also be worth of further research.

The study of the affinity-relevant correlations obtained with ANTIPASTI reveals long-range effects that are leveraged by the model to predict binding affinity, consistent with antigen-distal mutations being beneficial to affinity [52]. Similarly, non-additive dependencies (epistasis) between distant sites have been involved in changes in binding affinity upon joint mutations [64], with these dependencies stretching along all the CDRs and FRs of the heavy chain [30] and across the heavy and light chain [38]. These observations are also in line with the extended inter-region and inter-chain synergy in binding affinity suggested by our model. Formulating the prediction of binding affinity in terms of pairwise residue correlations through ANTIPASTI also offers a natural avenue for quantifying the synergy between regions, while avoiding the computationally costly evaluation of pairwise interactions through randomisation approaches. The synergistic patterns detectable in this way could be further explored in future as a help towards rational processes of acquisition of high binding affinity and specificity in antibodies. However, more data on the systematic assessment of the roles of residues in binding affinity by point or pairwise mutations with accompanying structures would be needed to clarify the mechanisms underlying affinity-relevant correlations uncovered by the ANTIPASTI learnt representations. We hope that the tools we provide here can contribute towards the goal of antibody design in conjunction with laboratory-controlled mutagenesis aimed at antibody affinity improvement. In this regard, a main challenge for the future would be to build upon these insights to develop a strategy to propose mutations that can lead to viable antibody structures with enhanced binding affinity.

## Acknowledgements

K.M. acknowledges support from the President’s PhD Scholarship at Imperial College London. M.B. acknowledges support by the Engineering and Physical Sciences Research Council (EPSRC) under grant EP/N014529/1 funding the EPSRC Centre for Mathematics of Precision Healthcare at Imperial College London. All authors are grateful to Francesco A. Aprile and Clément Nizak for their valuable feedback and suggestions.

## Author contributions

Study concept and design: K.M., M.B., and B.B.; Development of source code: K.M.; Analysis and interpretation of data: K.M., M.B., B.B.; Writing and revision of the manuscript: K.M., M.B., and B.B.; Study supervision: M.B. and B.B.

## Methods

### Data collection and pre-processing

The Structural Antibody Database (SAbDab) [12, 13] contains all the antibody structures available in the Protein Data Bank archive (PDB)^1^. We downloaded all bound antibody structures with affinity data from SAbDab as of 23 June 2023 in PDB format using the Chothia numbering [65].

Entries without affinity or antigen data, nanobodies consisting only of a light chain and single-chain variable fragments (scFvs) were discarded. Structures with less than three atoms in any residue, with a resolution worse than 4Å [66] or with very few residues in the heavy chain (less than 30) were excluded as well. Just 80 out of the remaining 634 antibodies (available in the GitHub repository) are nanobodies consisting of a heavy chain only, meaning that most of them have two paired chains. For both chains, we considered all amino acids in the variable region, corresponding, in Chothia numbering, to the range of positions from 1 to 113 for the heavy chain and from 1 to 107 for the light chain (Figure 1b).

### Normal Mode Analysis and the computation of correlation maps

The main goal of Normal Mode Analysis (NMA) is the characterisation of dynamical fluctuations around equilibria in physical systems [41]. This analysis is only valid near the equilibrium (*i.e*., for small deformations) and for short times. Still, Normal Modes have been useful for protein structures beyond this strict regime of validity, since they capture global information on the expected dynamical behaviour upon a perturbation (like the binding of a ligand at a certain site) that is a consequence of the connectivity of the protein’s network of residues. The key step of NMA is to simplify the dynamics using a set of generalised coordinates known as Normal Modes, which are derived from the spectral decomposition of the Hessian of the potential describing the system, as follows.

Let **r**^∗^ ∈ ℝ ^3*N*^ be the coordinates of a system of *N* particles in 3D space at equilibrium. Then, its potential energy under a small perturbation can be written (to second order) as:

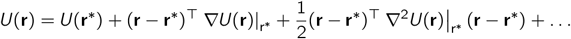

Here, *U*(**r**^∗^) is arbitrary and can be set to zero [67], while the gradient term is zero at a local minimum. Therefore, the potential energy can be expressed as a function of components of the Hessian matrix ∇^2^*U*(**r**) evaluated at the minimum **r**^∗^.

The solutions to the equations of motion under a quadratic potential as given by *U* take the form:

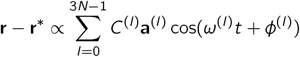

The fluctuations of the particles can be thus expressed as linear combinations of Normal Modes, indexed by *l*. Hence *C*^(*l*)^ and *φ*^(*l*)^ are, respectively, the am( plitude)and the phase of the *l* ^*th*^ Normal Mode, **a**^(*l*)^ ∈ ℝ^3*N*^ is the *l* ^*th*^ eigenvector of the Hessian matrix ∇^2^*U*(**r**)^∗^ and *ω*^(*l*)^ is its associated eigenvalue [68]. It is important to note that the first six NM are trivial, as they are associated to rotational and translational invariances [69].

A potential that assumes that each particle behaves as a harmonic oscillator [70] was proposed in Ref. [42] and underpins the so-called Elastic Network Models (ENMs):

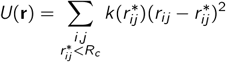

with *r*_*ij*_ the distance between particles *i* and *j, R*_*c*_ a cut-off radius and 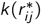 the pairwise force terms, here specified by the *α*-Carbon force field. The latter, derived from the fitting to the Amber94 [71] potential, is given by:

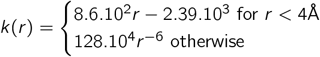

We use Bio3D [45] as an implementation of this formalism at the residue level to compute the Normal Modes of antibody-antigen complexes, and the associated Normal Mode correlation maps:

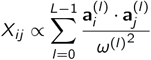

where is the dot product in ℝ^3^, 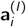 represents the Cartesian coordinates of the *l* ^*th*^ eigenvector of the Hessian matrix ∇^2^*U*(**r**) ^∗^ corresponding to residue *i*, and similarly for 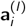. *L* is the number of modes considered, which can be at most 3*N*. The maps are then normalised to the [−1, 1] range.

In our work, we calculate these maps in the presence of the antigens to capture the imprint of the antigen correlations with the antibodies; once computed, the antigen residues are eliminated from the input maps for our algorithm. Missing residues (gaps) are encoded as zeros in the Normal Mode maps, denoting absence of correlation of their Normal Modes. The final value for *N* is 292 and it corresponds to the length of heavy and light chains with alignment to Chothia. This is performed identically for the nanobodies, but as expected the input maps are zero-valued in all pixels of the light chain and of the blocks shared by the heavy and light chains. To construct the NM correlation maps, we rely on the software [45], a package to compute the Normal Modes of a given protein structure, including fluctuations, deformation, and residue cross-correlation analysis.

### ANTIPASTI Neural Network architecture

ANTIPASTI is based on a convolutional neural network which receives as input Normal Mode correlation maps *X* ∈ ℝ^*N*×*N*^ (see Figure 1c). The model is PyTorch-based [72] and comprises a bank of *n*_f_ *k* × *k* convolutional filters, followed by a ReLU activation function and a *p* × *p* max pooling layer with stride *s*. The latter is followed by a bias-free fully-connected layer from which the value of log_10_(*K*_*D*_) is computed as follows. Let **z** ∈ ℝ^*M*^ be the flattened input of the ANTIPASTI fully-connected layer, produced by the network’s convolutional layer (with *n*_f_ filters and the non-linear activation) and the max pooling layer applied to a Normal Mode correlation map *X* ∈ ℝ^*N*×*N*^.

Let **w** ∈ ℝ^*M*^ be the weights of the fully-connected layer. Then, since there is no bias, the prediction of *K*_*D*_ is simply

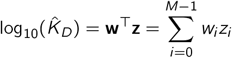

where for the described architecture

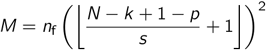

as every convolution operation yields a matrix of shape (*N* − *k* + 1) × (*N* − *k* + 1) which is subtracted by *p* and downsized by *s* through max pooling with stride *s* and kernel shape *p* × *p*. In this work, we take *s* always equal to *p*.

### Training and evaluation

We split 634 available correlation maps with their corresponding log_10_(*K*_*D*_) values into a training set and a test set in a 95 − 5 ratio. Training is performed in a supervised fashion with the AdaBelief optimiser and a learning rate *l*_*r*_. AdaBelief [73] guarantees the fast convergence found in adaptive methods such as Adam, good generalisation as in accelerated schemes such as Stochastic Gradient Descent (SGD) and non-exploding gradients during training. The parameters were set to *β*_1_ = 0.9, *β*_2_ = 0.999 and *ϵ* = 10^−8^. The latter is a typical configuration for a regression task [74].

Given a set of parameters *θ* = {*n*_f_, *k, p*}, a Lagrange multiplier *λ* ∈ ℝ_*>*0_, a batch of size *B*_*s*_ of input data points *X*^(*i*)^, and a function *f*_*θ*_ : ℝ^*N*×*N*^ → ℝ, the loss function used to learn *θ* is given by:

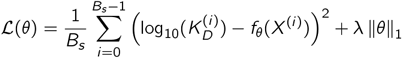

It consists of a data fidelity term and a regularisation term. The first one quantifies the mean squared error (MSE) between the ground truth log_10_(*K*_*D*_) and the prediction log_10_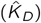 = *f*_*θ*_(*X*), and its expected value is computed as the empirical mean over a batch (*B*_*s*_ = 32). To favour sparsity in the weights of the neural network while avoiding overfitting, the *𝓁*_1_ regularisation [75] is employed in the second term with *λ* = 2*e* − 3.

Hyperparametric search was performed with Optuna [76] and 10-fold cross-validation on the training set by considering all the possible combinations of *n*_f_ ∈ {2, 4, 8}, *k* ∈ {3, 4, 5}, *p* ∈ {1, 2, 3} and *l*_*r*_ ∈ {5*e* − 5, 1*e* − 4, 5*e* − 4, 1*e* − 3}. With comparable accuracy, the top two candidates were (Figure S1):

- *n*_f_ = 4, *k* = 4, *p* = 2, and *l*_*r*_ = 1*e* − 4 (83000 parameters);
- *n*_f_ = 4, *k* = 4, *p* = 1, and *l*_*r*_ = 5*e* − 5 (334000 parameters).

The training was conducted for 5 different random seeded train-test splits using these optimal hyperparameters. The performance of ANTIPASTI was then evaluated on the test sets through the Pearson’s correlation coefficient *R* between predicted and ground truth binding affinity values (Figure 2a,c).

A linear regression model with no bias term and the same loss function was also considered. In essence, this model maps the input pixels to log_10_ (*K*_*D*_) using *N*^2^ weights.

Contact maps (with *α*-Carbon cutoff distances of 8Å [77]) were also examined for comparison with the Normal Mode correlation maps. The optimal hyperparameters were the same as for NMA, except that for the case without pooling the optimal learning rate *l*_*r*_ turned out to be 1*e* − 4 instead of 5*e* − 5.

### Interpretation of the learnt output layer representations

We construct 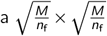 array *F* using the weights of the fully-connected layer **w** ∈ ℝ^*M*^ and its input **z** ∈ ℝ^*M*^. We refer to *F* as the *output layer representation* and it is obtained by reshaping the *M* output pixels of the fully-connected layer:

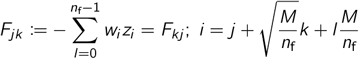

with *j* = 0, …, 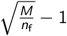; *k* = 0, …, 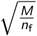 −1, then

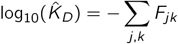

The positive (respectively, negative) pixels of *F* can be interpreted as the Normal Mode correlations in *X* responsible for its high (respectively, low) binding affinity. This is illustrated in Figure 5.

**Figure 5:**
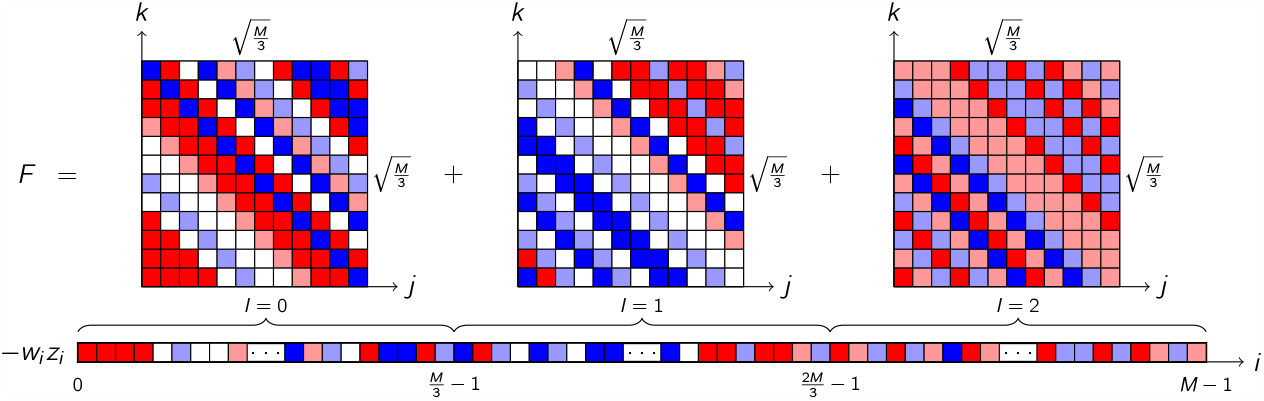
Construction of the *F* array for *n*_f_ = 3.

For the sake of better visualisation and inspection, the *F* arrays are post-processed with UMAP [51] using a minimum Euclidean distance of 0.1 and a minimum of 90 neighbours. We coloured the data points according to sequence properties extracted directly from SAbDab, such as antigen type (Figure 3b), V gene family, light chain type and antibody species (Figures S3a-c, Figures S2b-d). For the latter two, ellipses comprising 75% of the data points were added to show the separation of the different classes.

### Antibody regions importance

To study which heavy and light chain regions have a greater impact on the binding affinity, we computed the importance *I* for each region *s*^∗^ with a mean number of residues across the entire dataset 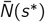 as follows:

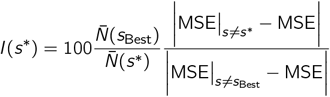

We estimated the importance *I*(*s*^∗^) separately for each group of antibodies binding to different antigen types (Figure 3c). The Mean Squared Error (MSE) was computed as:

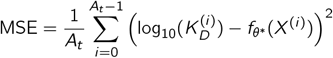

where *f*_*θ*∗_ denotes the binding affinity prediction log_10_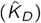 by the final ANTIPASTI architecture with learnt parameters *θ*^∗^. *A*_*t*_ is the number of antibodies in the training set binding to a given antigen type *t*, the types being proteins, peptides, haptens and carbohydrates (*A*_*t*_ = 347, 113, 53 and 24 respectively). Nanobodies were excluded as they would introduce a bias in the importance in favour of the heavy chain regions.

Excluding a region means that all output layer pixels in the region and their correlations with other regions are removed for the predictions. *s*_Best_ is the region that, when excluded, causes the MSE to deviate the most from that obtained with all regions. Hence *s*_Best_ represents the region that contributes the most to an accurate prediction of *K*_*D*_. Regions importance is defined in such a way as to be 100 for *s*_Best_, thus for each region it can be expressed as a percentage of *s*_Best_ importance.

We estimated the regions importance for the entire Framework and CDR regions of the heavy and light chains (Figure 3c), and at amino acid resolution (Figure S4b). In the latter case, each *s*^∗^ is an individual amino acid and 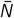(*s*^∗^) = 1 for all of them. To build Figure 3c, we examined for each region *s*^∗^ the proportion of MSE deviation attributable to correlations between residues within the same region (intra-region) and that resulting from correlations with residues in other regions (inter-region). We proceeded with the same logic for Figure S4a, this time considering correlations within the chain (heavy or light) to which *s*^∗^ belongs, as opposed to those established between *s*^∗^ and residues from the other chain.

### AlphaFold prediction

We used the ColabFold v1.5.2 [78] implementation of AlphaFold2. The antigen and antibody were chained together in one sequence and, for each prediction, 20 recycles with an early stop tolerance of 0.5 were used. We gave the possibility of detecting templates from a 100% clustered PDB, *i.e*., the option `template_mode` was set to `pdb100`. While retaining the original model training weights, ColabFold features a faster search via MMseqs2 [79] instead of using the time-consuming MSA generation step [80].

We used the 21 samples of the test set from one random training-test partition to make the comparison between the log_10_(*K*_*D*_) prediction of the original structure and that of the AlphaFold-estimated structure (Figure 4a). We refer to the subtraction of these two quantities as Δlog_10_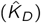 (Figure 4c). Specifically, the resulting PDB codes were `1a4k`, `1m7i`, `1tzi`, `1yej`, `2i5t`, `2p45`, `2r56`, `3bpc`, `3eoa`, `3o6l`, `3vw3`, `4f3f`, `4kht`, `4r8w`, `4rgo`, `4u6v`, `5cjq`, `5vzy`, `5w0k`, `5w1m` and `6axk`. Their labelled − log_10_(*K*_*D*_) values range between 4.8 and 11, hence are representative of a wide range of binding affinities.

When studying the relationship of the AlphaFold confidence metrics with Δlog_10_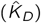 (Figure 4c), nanobody `2p45` appears as an outlier, in agreement with the observation that different ranges of AlphaFold prediction errors are associated to nanobodies [80]. Hence, it was excluded from the calculation of the correlation coefficients.

## Code availability

All the code is available in a GitHub repository as a Python package at github.com/kevinmicha/ANTIPASTI. It includes installation and dependency management instructions, ready-to-run environments and four tutorials in the notebook format. We also provide unit tests with 98.5% coverage of the total lines of code.

We wrote the code in a modular way, in such a way that it is possible to use NMA or contact maps, a CNN or LR and customise both models by just modifying the settings. This makes it immediately adaptable to architectures with more trainable parameters in the future if additional data becomes available.

## Footnotes

1 https://www.rcsb.org/

