## Supplemental information for "ANTIPASTI: interpretable prediction of antibody binding affinity exploiting Normal Modes and Deep Learning"

| Method | Based on | R score |
|---|---|---|
| Yang et al. (2023) [26] | Structure | 0.79 |
| Kurumida et al. (2020) [24] | Structure | 0.69 |
| Kang et al. (2021) [22] | Sequence | 0.66 |
| Sirin et al. (2015) [23] | Sequence | 0.45 |

**Table S1: Comparison to existing methods.** $R$ score reported for binding affinity prediction methods which, to the best of our knowledge, are the best performing in the literature.

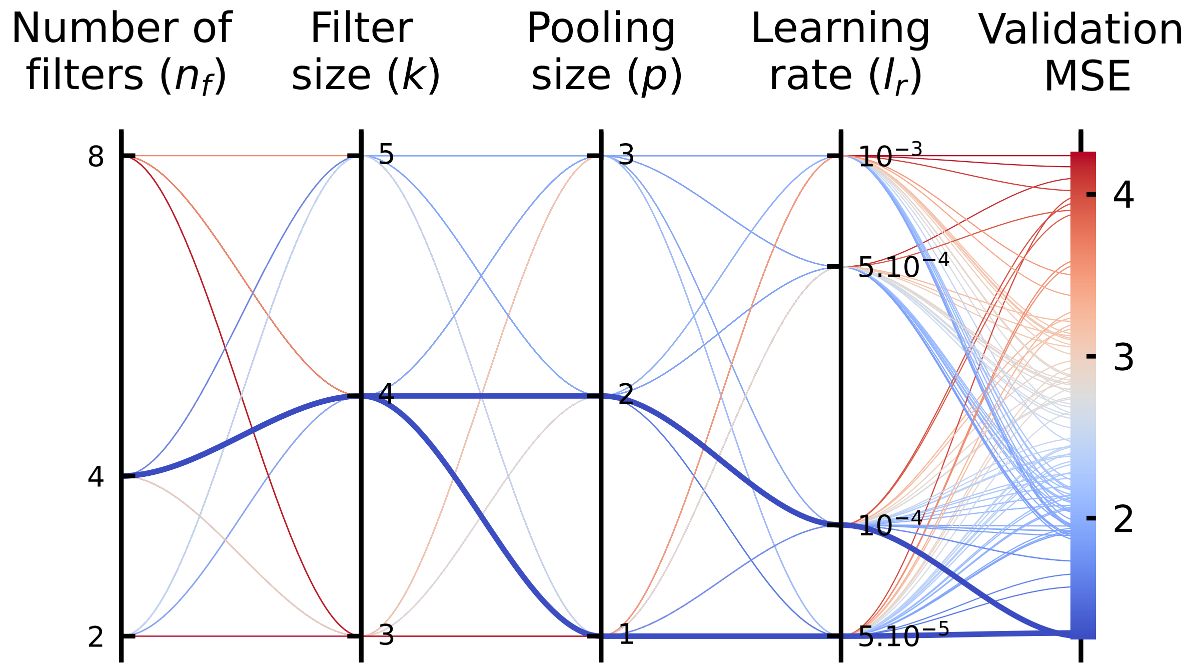


**Figure S1: ANTIPASTI model selection:** Optuna and 10-fold cross-validation. The two hyperparameter sets with the lowest validation MSE are represented by thicker lines and correspond to the two ANTIPASTI architectures whose performance is shown in Figures 2a-c (CNN with pooling $p = 2$ and without pooling).

### Antigen-agnostic case

We tested the scenario in which we train ANTIPASTI without taking antigen information into account when calculating the Normal Modes. We call this scenario the *antigen-agnostic* case, which requires a bound antibody-antigen conformation but not the antigen sequence or its atom coordinates. Although the accuracy (Figure S2a) is lower than that of the standard case (with *antigen imprint*), we found, when following the same procedure as for Figure 3b, noteworthy separations according to various biological properties in the UMAP maps of the output layer representations (Figures S2b–d).

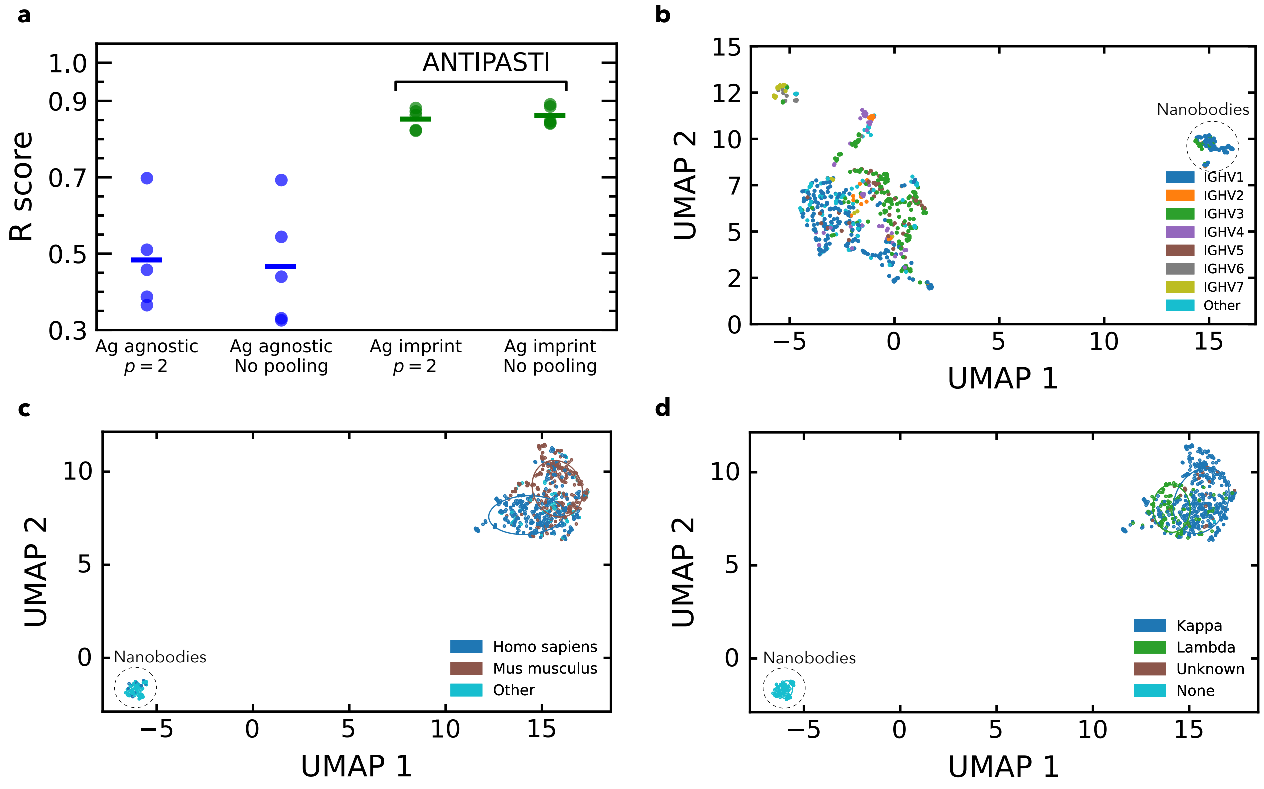


**Figure S2: The antigen-agnostic case**. (**a**) Same plot as in Figure 2c, but comparing the two top-performing models with their antigen-agnostic analogues. (**b**, **c**, **d**) UMAP of the output layer representations with data points using colour codes that correspond to: heavy chain V gene, antibody species, and type of light chain respectively.

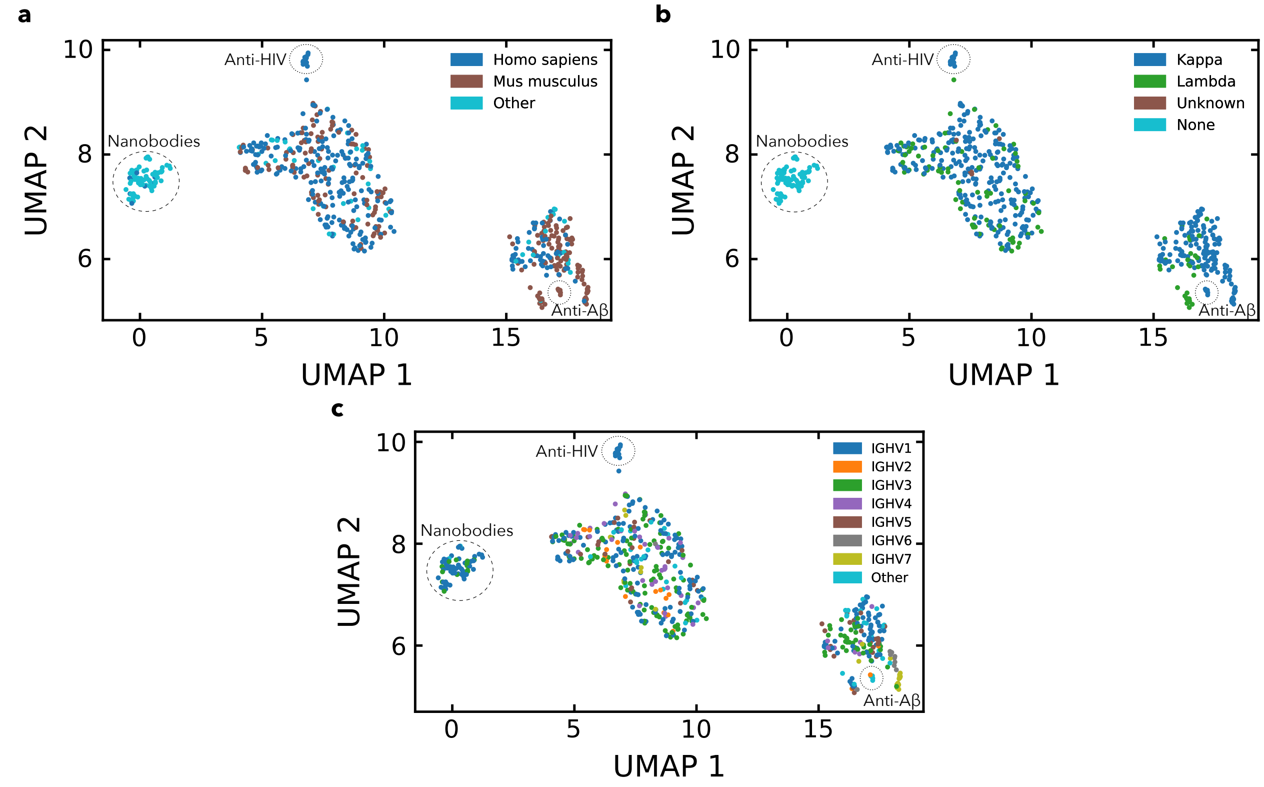


**Figure S3: UMAP of the output layer representations (with antigen imprint)**. Data points coloured according to: (**a**) Antibody species. (**b**) Type of light chain. (**c**) Heavy chain V gene.

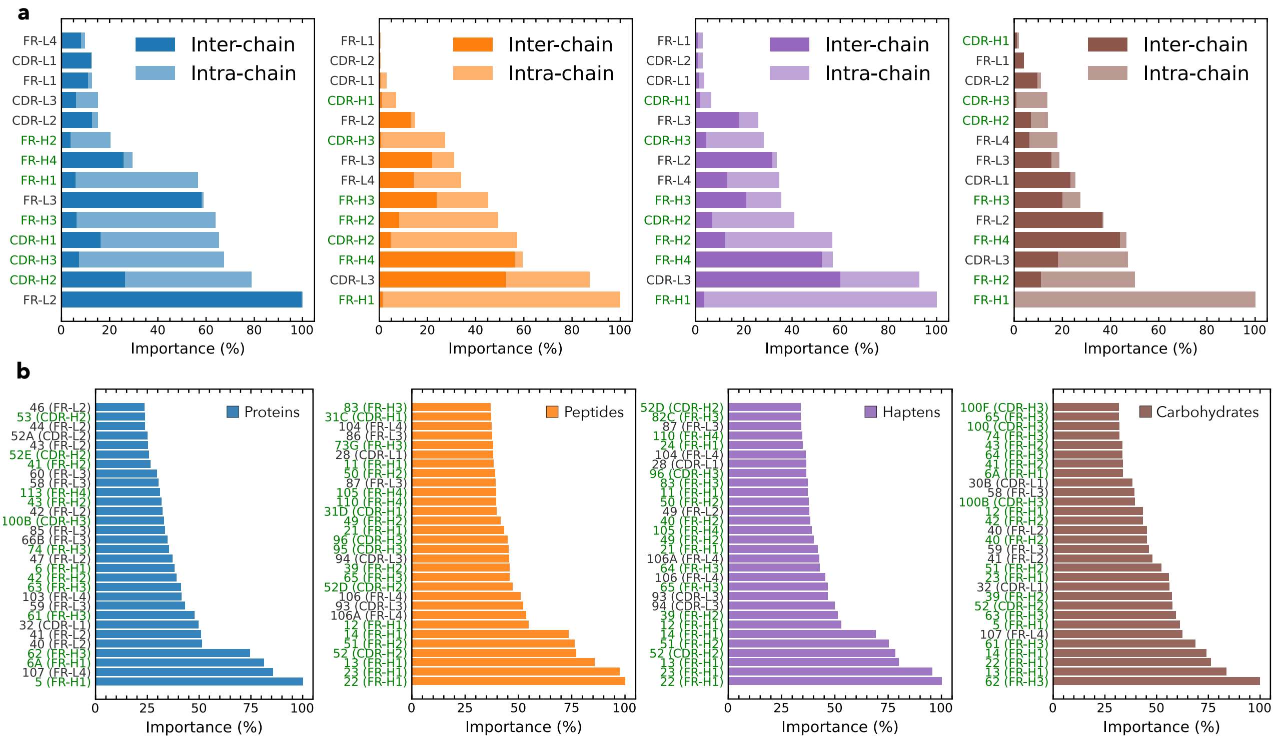


**Figure S4: Regions and residues importance**. (**a**) Proportion of MSE deviation attributable to correlations between residues within the same chain (intra-chain) and that resulting from correlations with residues in the other chain (inter-chain). (**b**) The ranking of importance of single amino acids varies depending on the type of target for antibodies with paired heavy and light chains.

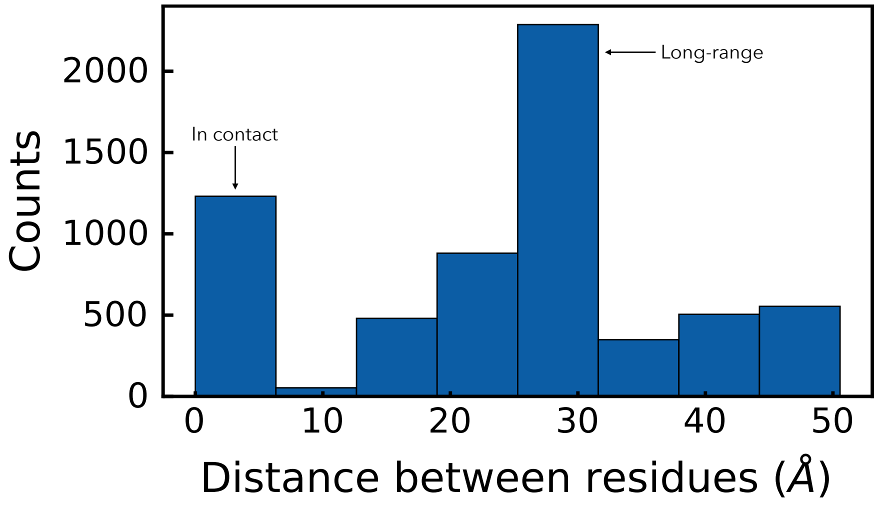


**Figure S5:** Distribution of residue $\alpha$-Carbon pairwise distances involved in the top 10 affinity-relevant correlations for binding affinity across all 634 antibody structures.

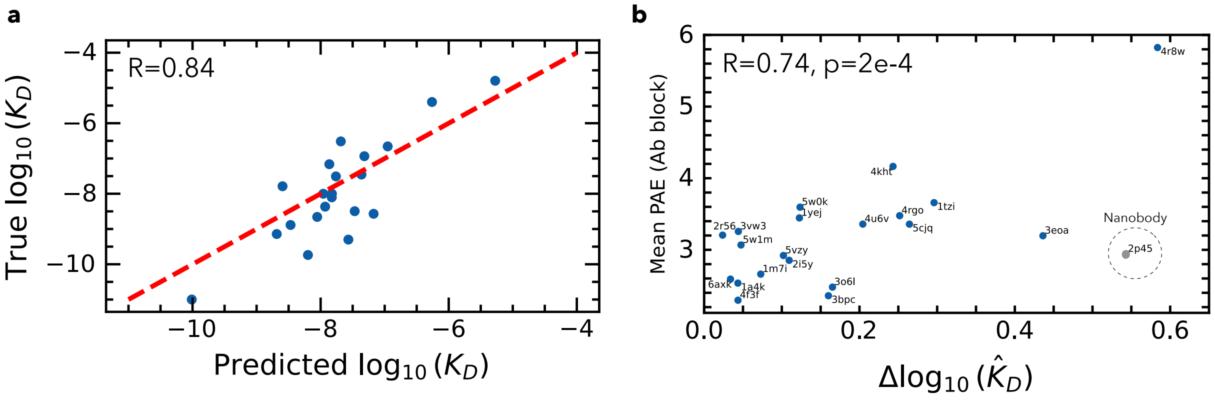


**Figure S6: AlphaFold-predicted structures**. (**a**) ANTIPASTI predictions for a particular test set that yielded a correlation coefficient $R = 0.84$. (**b**) Mean predicted alignment error (PAE) for the antibody residues only, as a function of $\Delta\log_{10}(\hat{K}_D)$.
